# Characterization of the Novel Mitochondrial Genome Replication Factor MiRF172 in *Trypanosoma brucei*

**DOI:** 10.1101/194811

**Authors:** Simona Amodeo, Martin Jakob, Torsten Ochsenreiter

## Abstract

The unicellular parasite *Trypanosoma brucei* harbors one individual mitochondrial organelle with a singular genome the kinetoplast DNA or kDNA. The kDNA largely consists of concatenated minicircles and a few maxicircles that are also interlocked into the kDNA disc. More than 30 proteins involved in kDNA replication have been described, however several mechanistic questions are only poorly understood. Here, we describe and characterize MiRF172, a novel mitochondrial genome replication factor, which is essential for proper cell growth and kDNA maintenance. Using super-resolution microscopy, we localize MiRF172 to the antipodal sites of the kDNA. We demonstrate that depletion of MiRF172 leads to continuous loss of mini- and maxicircles during the cell division cycle. Detailed analysis suggests that MiRF172 is likely involved in the reattachment of replicated minicircles to the kDNA disc. Furthermore, we provide evidence that the localization of the replication factor MiRF172 not only depends on the kDNA itself, but also on the mitochondrial genome segregation machinery suggesting a tight interaction between the two essential entities.

**Summary Statement:** MiRF172 is a novel protein involved in the reattachment of replicated minicircles in *Trypanosoma brucei*, which requires the mitochondrial segregation machinery for proper localization.

## Introduction

One of the most intriguing genome organizations can be found in the mitochondrial genome of Kinetoplastea a class of single celled eukaryotes. The name Kinetoplastea refers to the organism’s single mitochondrial genome (kinetoplast DNA, kDNA) that in most cases is positioned close the base of the flagellum. The position reflects the physical connection between the base of the flagellum and the kDNA by a structure called the tripartite attachment complex (TAC) as has been demonstrated in *Trypanosoma brucei* (Ogbadoyi, 2003). Several components of this structure have now been identified that elute to a rather complex organization of the segregation machinery (Gheiratmand et al., 2013; Käser et al., 2016; Käser et al., 2017; Schnarwiler et al., 2014; Trikin et al., 2016; Zhao et al., 2008). The mitochondrial genome itself is composed of small and larger plasmid like elements referred to as the mini- and maxicircles, respectively. Maxicircles (23 kb, in *T. brucei*) are the functional homologues of other mitochondrial genomes and encode 18 protein coding genes and two ribosomal RNA genes. The majority of the mitochondrial genes are cryptic and require posttranscriptional editing to code for the *bona fide* components of the respiratory chain and a ribosomal protein (Hajduk and Ochsenreiter, 2010; Jensen and Englund, 2012; Povelones, 2014). The editing process is mediated by large protein complexes (Aphasizheva and Aphasizhev, 2016; Göringer, 2012; McDermott et al., 2016) and small non-coding guide RNAs (gRNAs) that are transcribed from the minicircles, which in *T. bruce*i are 1 kilo base (kb) in size and each of them code for three to five gRNAs (Hajduk and Ochsenreiter, 2010; Hong and Simpson, 2003; Ochsenreiter et al., 2007). Each *T. brucei* cell contains a single mitochondrion with one kDNA. The kDNA is made up of 5000 minicircles with several hundred different minicircle classes, and 25 maxicircles (23 kb), which are virtually identical. Each minicircle is physically connected to three other minicircles and the maxicircles are interwoven in the minicircle network (Chen et al., 1995). Overall the kDNA resembles a chain mail and is organized likely through several histone like proteins (Lukeš et al., 2001; Xu et al., 1996) in a disc like structure about 450nm in diameter and 150nm in height (Jakob et al., 2016). Similar to the overall structure, replication of the kDNA is complex and some estimate up to 150 proteins to be involved in this process (Jensen and Englund, 2012). The current replication model suggests that the minicircles are released from the network into the kinetoflagellar zone (KFZ), the region between the kinetoplast and the inner mitochondrial membrane (Drewa and Englunda, 2001; Jensen and Englund, 2012). Here replication is initiated and proceeds unidirectionally via theta intermediates leading to two nicked / gapped minicircles that are then transported to the antipodal sites via an unknown mechanism (Jensen and Englund, 2012; Povelones, 2014). The antipodal sites are ill defined protein complexes at opposing sites of the kDNA disc. Within these sites partial gap repair occurs and the minicircles are reattached to the kDNA network by a Topoisomerase type II enzyme (Wang et al., 2000). The newly replicated and reattached minicircles maintain at least one gap each until the networks are separated. The repair of the gaps (and nicks) is likely mediated by the mitochondrial ligase LigK alpha (Downey et al., 2005) and the mitochondrial polymerase Pol beta PAK (Saxowsky et al., 2003). After, the duplicated kDNA is segregated by the separating basal bodies that are connected to the kDNA via the TAC structure (Ogbadoyi, 2003). Several core components of the three regions of the TAC have now been identified: p197 and TAC65 in the exclusion zone filaments (EZF) (Gheiratmand et al., 2013; Käser et al., 2016), TAC40 in the outer mitochondrial membrane (Schnarwiler et al., 2014), p166 at the inner mitochondrial membrane (Zhao et al., 2008) and TAC102 in the unilateral filaments (ULF) (Trikin et al., 2016). We have now also elucidated the hierarchy of the TAC complex and its assembly demonstrating that the structure is built *de novo* from the basal body towards the kDNA and that TAC102 is currently the kDNA most proximal TAC protein, while p197 is closest to the basal body (Hoffmann et al., 2017).

Here we present data characterizing the **Mi**nicircle **R**eplication **F**actor **172** (MiRF172; Tb927.3.2050) as a kDNA associated protein essential for proper growth and kDNA maintenance in *T. brucei*. We also demonstrate that MiRF172 is likely involved in the reattachment of replicated minicircles to the kDNA disc. Further we demonstrate that localization of MiRF172 partially depends on the TAC and not only on the kDNA, suggesting a tight interaction between replication and segregation machinery.

## Results

MiRF172 is a hypothetical conserved very basic (pI 9.5), large (172 kDa) protein with a predicted mitochondrial targeting sequence at the N-terminus (Fig. 1B). MirRF172 has been detected in several proteomics studies (i) supporting its mitochondrial localization (Peikert et al., 2017; Zhang et al., 2010), (ii) showing its developmental regulation (Gunasekera et al., 2012) and (iii) indicating a possible phosphorylation site at S999 (Urbaniak et al., 2013). The gene including its position in the genome is conserved throughout the currently sequenced Kinetoplastea (Fig. 1A). The protein contains a poly-Q stretch and an alanine-lysine rich region, both of which are found in the C-terminal part (Fig. 1B). While the poly-Q stretch is only conserved among the Trypanosoma species, the alanine-lysine rich region is conserved throughout the Kinetoplastea.

**Fig. 1.**
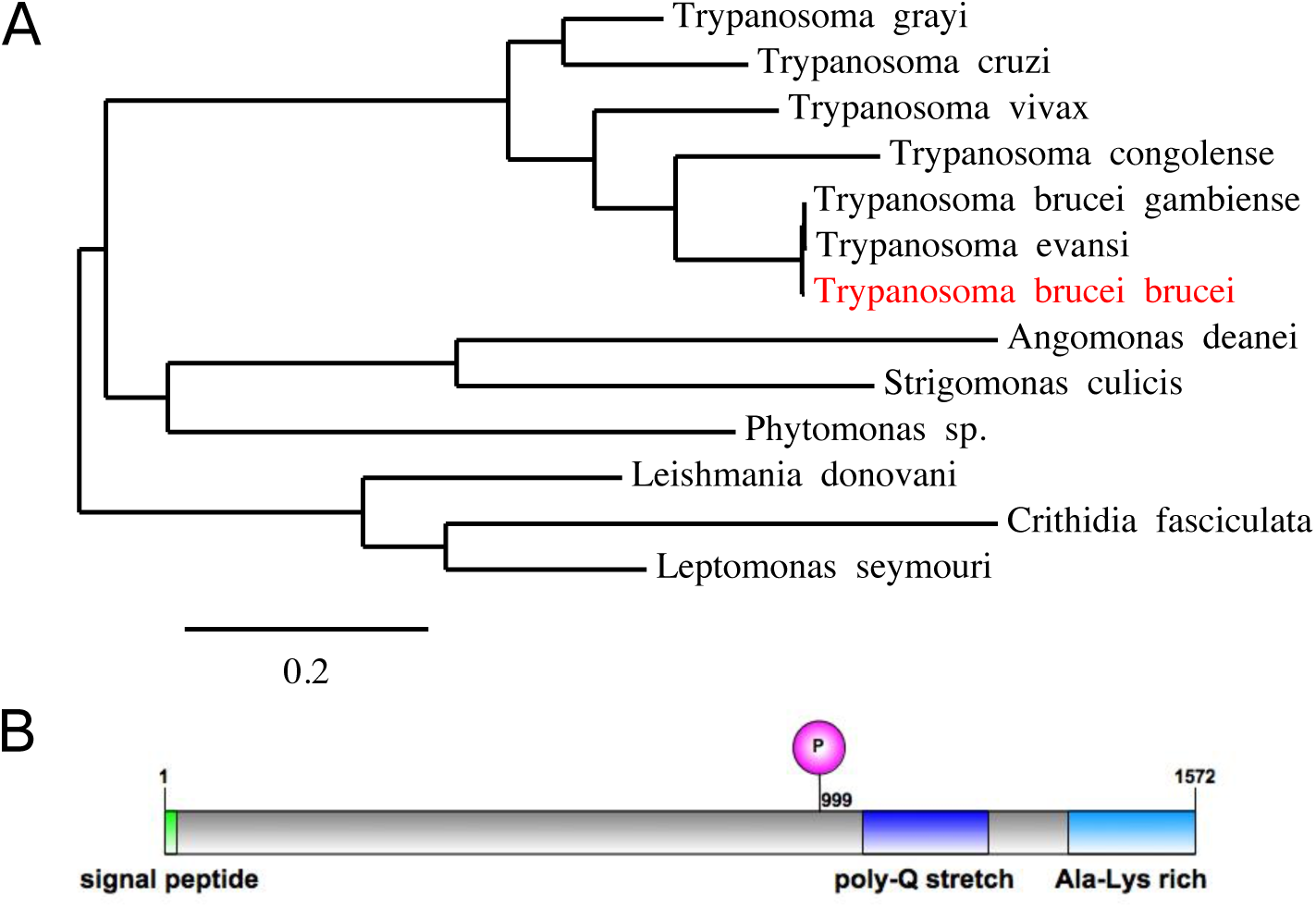
Protein properties of MiRF172 in *T. brucei* cells. **A)** A phylogenic tree showing the conservation of MiRF172 among Kinetoplastids. MiRF172 is highlighted in red. **B)** Illustration of MiRF172 ORF. Depicted are in green the signal peptide for mitochondrial import, in magenta the phosphorylation site at position 999, in dark blue the poly-Q stretch enriched domain and in light blue the alanine and lysine enriched C-terminal domain.

### MiRF172 protein localizes at the kDNA

To localize the MiRF172 protein, we tagged it *in situ* at the C-terminus with a PTP epitope tag in blood stream form (BSF) and with HA in procyclic form (PCF) *T. brucei* (Fig. 2). Based on colocalization studies in BSF cells with the basal body marker YL1/2 and the DNA stain DAPI the protein localizes between the basal body and the kDNA in the KFZ (Fig. 2A). MiRF172 is expressed throughout the cell cycle in both lifecycle stages (Fig. 2B, C). The protein forms two foci 180° apart on kDNA discs in 1K1N, 2K1N and 2K2N (K = kinetoplast, N = nucleus, 1K1N = cells are in G1 of the cell cycle, 2K1N= cells are in nuclear S phase, 2K2N = cells just prior to cytokinesis) cells in both life cycle stages (Fig. 2B, C; Fig. 3A upper panel, 93% of the cells showed this MiRF172 localization) reminiscent of the antipodal sites that have been described for many kDNA associated proteins (Jensen and Englund, 2012). In rare cases, we also observed localization of MiRF172 covering the whole disk (in 1% of 1K1N, 2K1N or 2K2N cells) or forming circles around the whole kDNA disk (in 1% of 1K1N, 2K1N or 2K2N cells). During replication of the kDNA when the mitochondrial genome adopts a bilobed structure (Fig. 2B, C; d1K1N) MiRF172 remains as two foci on the opposing sites (in 64% of all d1K1N, Fig. 3A lower panel) until just prior to the division of the kDNA, when a third spot appears in the middle between the two segregating discs (in 36% of all d1K1N cells, Fig. 3A lower panel, model Fig. 3C). After segregation, the second spot is present on each of the kDNA disc (model Fig. 3C). In 3D reconstructions of 1K1N kDNA discs using super resolution STED imagery the protein forms two curved structures each covering about 25% of the kDNA circumference facing the KFZ (Fig. 3B). We also used biochemical approaches to isolate mitochondria as described previously (Trikin et al., 2016). Solubilization of the mitochondrial fraction with 1% digitonin leads to an insoluble and soluble fraction. MiRF172 remained mostly associated with the insoluble fraction even after DNAseI treatment (Fig. S2B, C). Furthermore, we isolated flagella from the cells as described previously (Ogbadoyi, 2003) and could show that MiRF172 remains associated with flagella (Fig. S2A) similar to what has been described for TAC components (Gheiratmand et al., 2013; Käser et al., 2016; Trikin et al., 2016; Zhao et al., 2008).

**Fig. 2.**
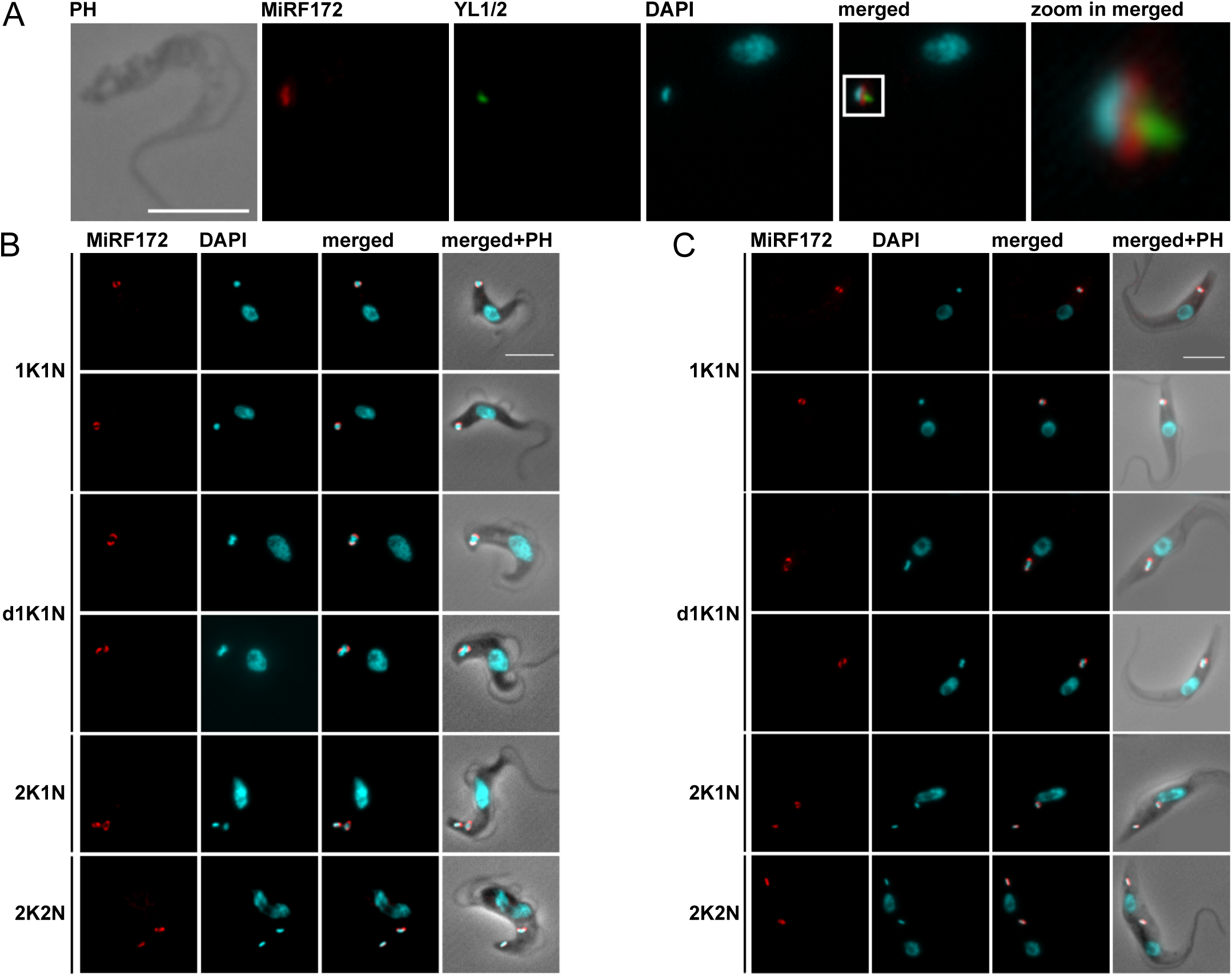
Localization of MiRF172 in BSF and PCF *T. brucei* cells. **A)** Immunofluorescence microscopy of MiRF172-PTP tagged BSF cells. Localization of MiRF172-PTP (red) is represented by maximum intensity projections from immunofluorescence microscopy image stacks of *T. brucei* BSF cells. MiRF172-PTP was detected by the α-Protein A antibody. The mature basal bodies were detected with the YL1/2 monoclonal antibody (green). The kDNA and the nucleus were stained with DAPI (cyan). **B)** Immunofluorescence analysis of MiRF172-PTP during different stages of the cell cycle (1K1N, dK1N, 2K1N, 2K2N) in BSF cells. K = kDNA, N = nucleus, dK = duplicating kDNA. Localization of MiRF172-PTP (red) and DNA (cyan) were performed as described in A. **C)** Immunofluorescence analysis of MiRF172-HA during different stages of the cell cycle (1K1N, dK1N, 2K1N, 2K2N) in PCF cells. Localization of MiRF172-HA (red) represented by maximum intensity projections from immunofluorescence microscopy image stacks of PCF cells. MiRF172-HA was detected by the α-HA antibody. The kDNA and the nucleus were stained as describe in A. PH = phase contrast. Scale bar = 5 μm

**Fig. 3.**
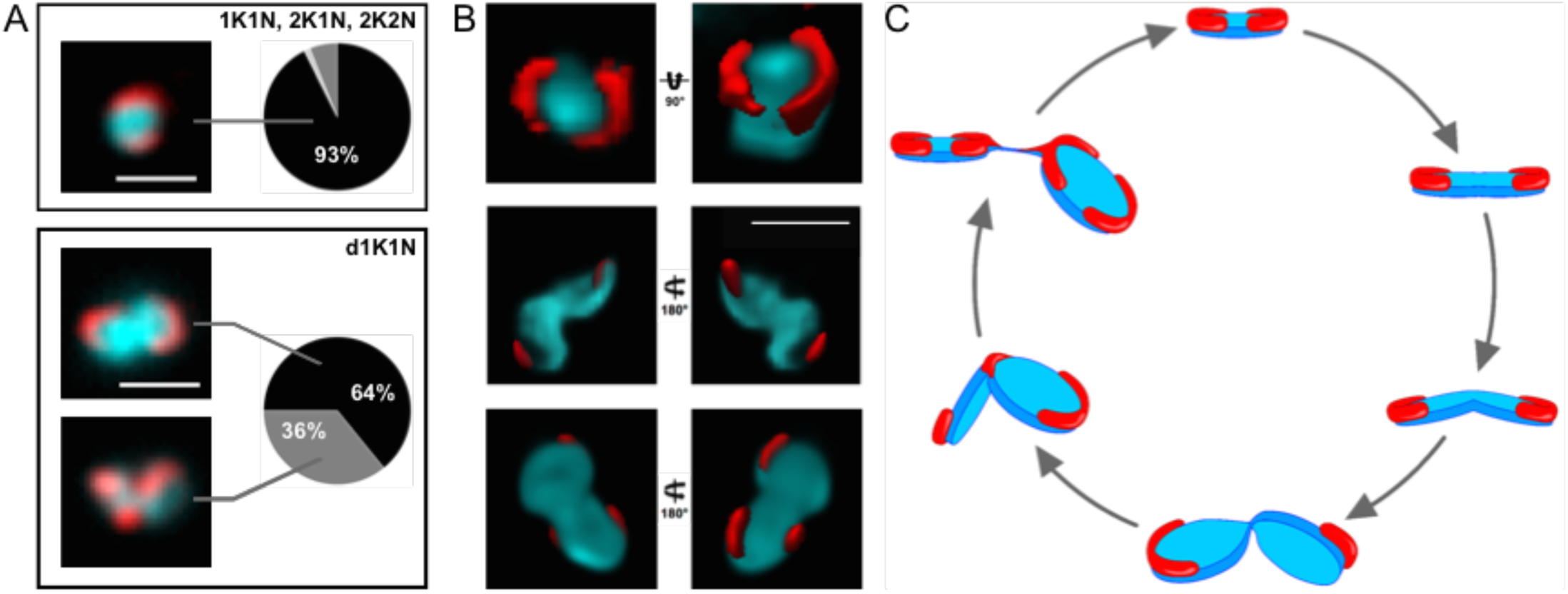
Analysis of MiRF172 localization during the cell cycle. **A)** Quantification of MiRF172-PTP localization at single or duplicated kDNAs (1K1N, 2K1N, 2K2N) and duplicating kDNAs (d1K1N) in BSF cells. K = kDNA, dK = duplicating kDNA, N = nucleus. The left side shows representative immunofluorescence microscopy images depicting the localization of MiRF172-PTP (red) relative to the kDNA disk (cyan). The pie charts show the localization of MiRF172 in the respective kDNA replication stage. In 93% of the 1K1N, 2K1N and 2K2N subpopulations, MiRF172 is located at the antipodal sites. **B)** 3D-STED immunofluorescence analysis of MiRF172-PTP in *T. brucei* BSF cells. MiRF172 (red) and kDNA (cyan) 3D projection (surface rendering) from different angles. MiRF172-PTP was detected by the α-Protein A antibody and acquired with 3D-STED. The kDNA was stained with DAPI (cyan) and acquired with confocal microscopy. Pictures were deconvolved with the Huygens professional software. **C)** Model of MiRF172 localization during the cell cycle. Depicted is a model of the different stages of kDNA disk (cyan) replication in *T. brucei* and the localization of MiRF172 (red) relative to the kDNA. Scale bars = 1 μm

### RNAi of MiRF172 leads to growth retardation and kDNA loss

To study the function of MiRF172 we depleted the mRNA by RNAi in NYsm BSF cells using the tetracycline inducible RNAi vector pTrypRNAiGate. Northern blot analysis showed a decrease of MiRF172 mRNA by 68% on day three of induction (Fig. 4A). After RNAi induction cells grow at normal rates until day four when a growth defect becomes visible that is maintained at least until day eight. The growth defect was not accompanied by any obvious change in cell morphology or motility. In order to characterize a potential effect of MiRF172 depletion on mitochondrial genome replication we sampled the population (n ≥ 100 for each condition and replicate) at day zero and three post induction, stained the cells with the DNA dye DAPI and evaluated the relative occurrence of kDNA and nucleus in different cell cycle stages: Cells with one kDNA and one nucleus (1K1N; cells are in G1 of the cell cycle), cells with already replicated and segregated kDNAs and one nucleus (2K1N, cells are in nuclear S phase), as well as cells that had replicated both, the kDNA and the nucleus (2K2N, cells just prior to cytokinesis). We also screened for any abnormal K-N combinations like 1K2N (likely a product of kDNA missegregation), 1K0N (zoid cells) as well as 0K1N (indicative of kDNA replication/segregation defects). The major change in K-N combinations was the accumulation of 0K1N cells to about 20% at day three post induction, just prior to the appearance of the growth phenotype (Fig. 4B, C). At the same time point we observed that 30% of 1K1N cells had smaller kDNAs than in the uninduced cells (Fig. 4B, C).

**Fig. 4.**
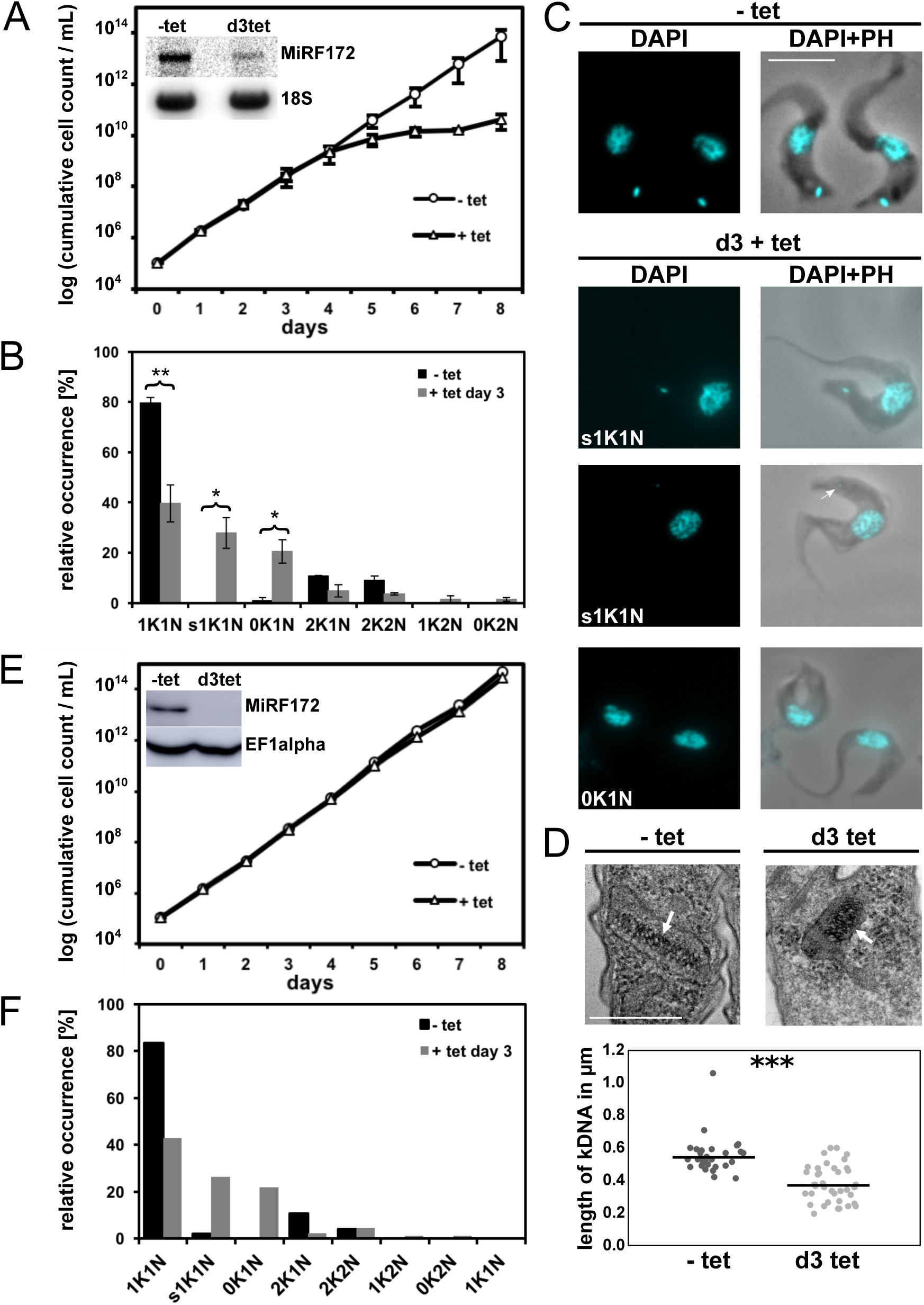
Phenotype upon knockdown of MiRF172 mRNA by RNAi in *T. brucei* BSF cells. **A)** Growth curve of MiRF172 RNAi *T. brucei* BSF cells. The y-axis shows the cumulative number of cells. Inset depicts a northern blot showing ablation of MiRF172 mRNA at day 3 post induction. 18S rRNA serves as a loading control. **B)** Quantification of the relative occurrence of kDNA and nucleus in MiRF172 RNAi induced and uninduced cells. K = kDNA, N = nucleus, sK = small kDNA. The y-axis shows the relative occurrence in the population. Significance of the differences was calculated by two-tailed unpaired t-test. * = p ≤ 0.05, ** = p ≤ 0.01. **C)** Representative fluorescence microscopy images of MiRF172 RNAi BSF cells. The nucleus and the kDNA were stained with DAPI. PH = phase contrast. Scale bar = 5 μm **D)** *Upper part*: Representative images of ultra-structures of the kDNA of MiRF172 RNAi cells revealed by TEM. Scale bar = 500 nm. *Lower part*: Length measurements of kDNA ultra-structures from uninduced and induced (three days) MiRF172 RNAi BSF cells. Y-axis shows length of kDNAs in microns. Significance of difference in length was calculated by two-tailed unpaired t-test. *** = p ≤ 0.001 **E)** Growth curve of MiRF172 RNAi BSF γL262P *T. brucei* cells. The inset depicts a western blot showing ablation of MiRF172-PTP protein at day 3 post induction. EF1alpha serves as a loading control. **F)** Quantification of the relative occurrence of kDNA and nucleus in MiRF172 RNAi γL262P *T. brucei* cells.

### RNAi of MiRF172 leads to an accumulation of smaller kDNA networks

To better characterize the kDNA loss phenotype we performed thin section transmission electron microscopy (TEM) and examined the ultra-structure of the kDNA networks of BSF cells induced for RNAi and compared them to uninduced cells. We measured the length of the striated structure that corresponds to a cross section through the kDNA disc in ≥ 30 randomly acquired kDNA images in each; uninduced and induced cells. In uninduced cells, the mean diameter of the kDNA was 544 nm while in induced cells it was significantly (p-value = 2.71×10^−8^) reduced to 368 nm (Fig. 4D, S1). Although the size (diameter) of the network was reduced, the overall appearance of the striated structure and its relative position to the basal body did not change.

### RNAi of MiRF172 in γL262P BSF cells has no influence on growth

In order to test if MiRF172 has essential functions that are not directly related to kDNA maintenance and whether the kDNA loss phenotype is a secondary effect, we used the recently established γL262P cell line that harbors a single point mutation in the F_1_F_0_-ATPase and is able to compensate for the loss of the kinetoplast similar to “petite” mutants in yeast (Dean et al., 2013). γL262P BSF cells were transfected with the inducible RNAi vector, which was previously used to generate the MiRF172 RNAi BSF cell line. We essentially performed the same analysis as described above and observed that, while the γL262P MiRF172 RNAi cells lost kDNA at a similar rate (n ≥ 100 for each condition) as the NYsm strain, the cells showed no additional growth phenotype suggesting that the sole function of MiRF172 is in kDNA maintenance and loss of kDNA is a direct effect of the depletion of MiRF172 (Fig. 4E, F).

### RNAi of MiRF172 leads to a loss of mini- and maxicircles

The results described above suggest that MiRF172 is involved in kDNA replication. To study the effect of kDNA loss in more detail, we performed Southern blot analyses of mini- and maxicircles in MiRF172 RNAi BSF cells. Whole cell DNA was extracted at day 0, day 3, day 5 and day 7 upon RNAi induction. The DNA samples were digested with HindIII and XbaI, resolved on an agarose gel and blotted on nylon membranes and probed for maxi- and minicircles. As a loading control, we used a probe targeting tubulin. We performed three biological replicates. Significance of the results was calculated using the two-tailed unpaired t-test (mini- and maxicircles d0 vs. d7 p ≤ 0.05 = *, covalently closed minicircles d0 vs. d5 p ≤ 0.01 = **, nicked / gapped minicircles p ≤ 0.05 =*). We detected a steady decrease of maxi- and minicircle abundance to about 60% of the uninduced levels from day zero to day five post induction of RNAi after which the amount of maxi- and minicircles increased again slightly (Fig. 5A, B). To further study the effect of MiRF172 depletion on minicircle replication we performed Southern blot analysis of minicircles released from the network, prior and post replication, respectively. For this whole DNA was extracted from uninduced and MiRF172 depleted cells at day 3, 5 and 7 upon RNAi. Southern blotting was done as described above but without restriction digest of the DNA (Fig. 5C, D). We detected a steady decrease of the covalently closed minicircles that have been released from the network but have not yet been replicated. For the nicked / gapped population, that represents the newly replicated intermediates prior to reattachment, increased significantly (p ≤ 0.05) until day five post induction. They then returned to the initial levels (Fig. 5C, D). Based on these results we suggest that MiRF172 is involved in the replication of the kDNA and more specifically in the reattachment process of the replicated minicircles to the kDNA network.

**Fig. 5.**
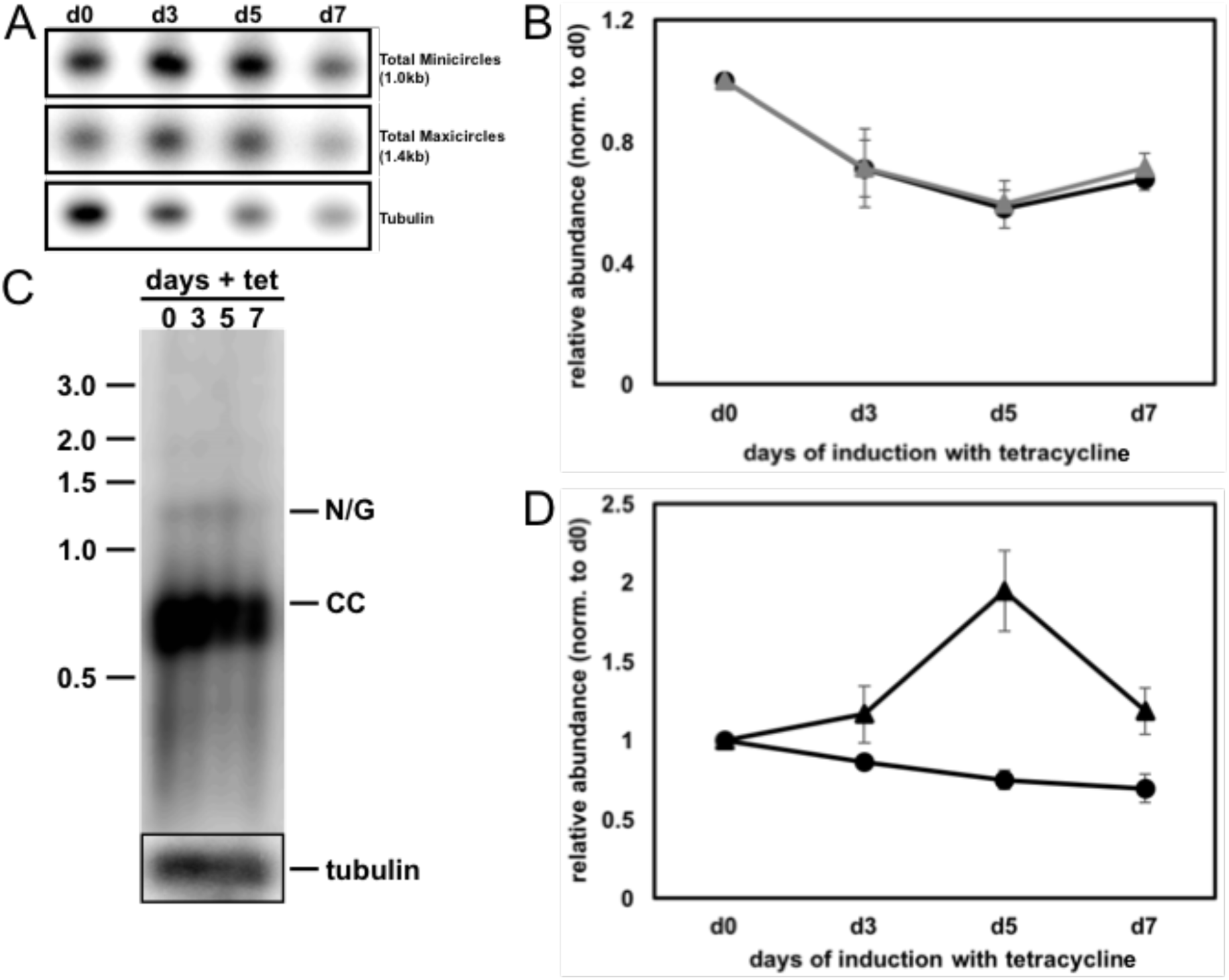
Effect of MiRF172 RNAi on kDNA abundance and free minicircle replication intermediates in *T. brucei* BSF cells. **A)** Detection of mini- and maxicircles by Southern blot. Total DNA (5 μg/lane) isolated from either uninduced (d0) or RNAi induced (d3, d5, d7) cells was digested with HindIII / XbaI, Southern blotted, and probed for minicircles (1.0-kb linearized minicircles), maxicircles (a 1.4-kb fragment), and tubulin as a loading control (a 3.6-kb fragment). **B)** Quantification of mini- and maxicircle abundance on Southern blots during MiRF172 depletion. Black circles = minicircles, grey triangles = maxicircles. Abundance ratio of total minicircle or maxicircle relative to the loading control (tubulin), normalized to day 0 of RNAi induction. Minicircles p-value d0 vs. d7 (p ≤ 0.05 = *), maxicircles p-value d0 vs. d7 (p ≤ 0.05 = *). **C)** Detection of free minicircle replication intermediates by Southern blot. Total DNA (5 μg/lane) isolated from either uninduced (d0) or RNAi induced (d3, d5, d7) cells, Southern blotted, and probed for minicircles. Nicked / gapped (N/G) and covalently closed (CC) minicircles are indicated. Tubulin was used as a loading control. **D)** Quantification of CC and N/G minicircles on Southern blot during MiRF172 depletion. Black circles = CC minicircles, black triangles = N/G minicircles. Abundance ratio of minicircles relative to the loading control tubulin and normalized to day 0 of RNAi induction. CC minicircles p-value d0 vs. d5 p ≤ 0.01 = **, N/G minicircles p-value d0 vs. d5 p ≤ 0.05 = *

### RNAi of MiRF172 has no impact on the TAC

Based on the observation that MiRF172 remains associated with the flagellum after flagellar extraction from BSF cells (same as TAC102, Fig. S2A) and that MiRF172 localizes in the region of the KFZ (Fig. 1A) we wondered if the protein would colocalize with a TAC marker protein of the ULF such as TAC102. For this we used immunofluorescence microscopy. The imagery shows that MiRF172 is located between TAC102 and the kDNA disc with little to no overlap between the two MiRF172 signals and the TAC102 signal (Figure 6A). As mentioned above biogenesis of the second signal for MiRF172 at the kDNA disc occurs during kDNA division and thus prior to the replication of the TAC102 signal (Hoffmann et al., 2017; Trikin et al., 2016). To test whether depletion of MiRF172 has an impact on TAC biogenesis, we probed for TAC102 during three days of MiRF172 RNAi. Even though we observed the typical MiRF172 depletion phenotype including increase in 0K1N cells and cells with smaller kDNAs, however we did not detect a loss in the signal for TAC102 (Fig. 6B).

**Fig. 6.**
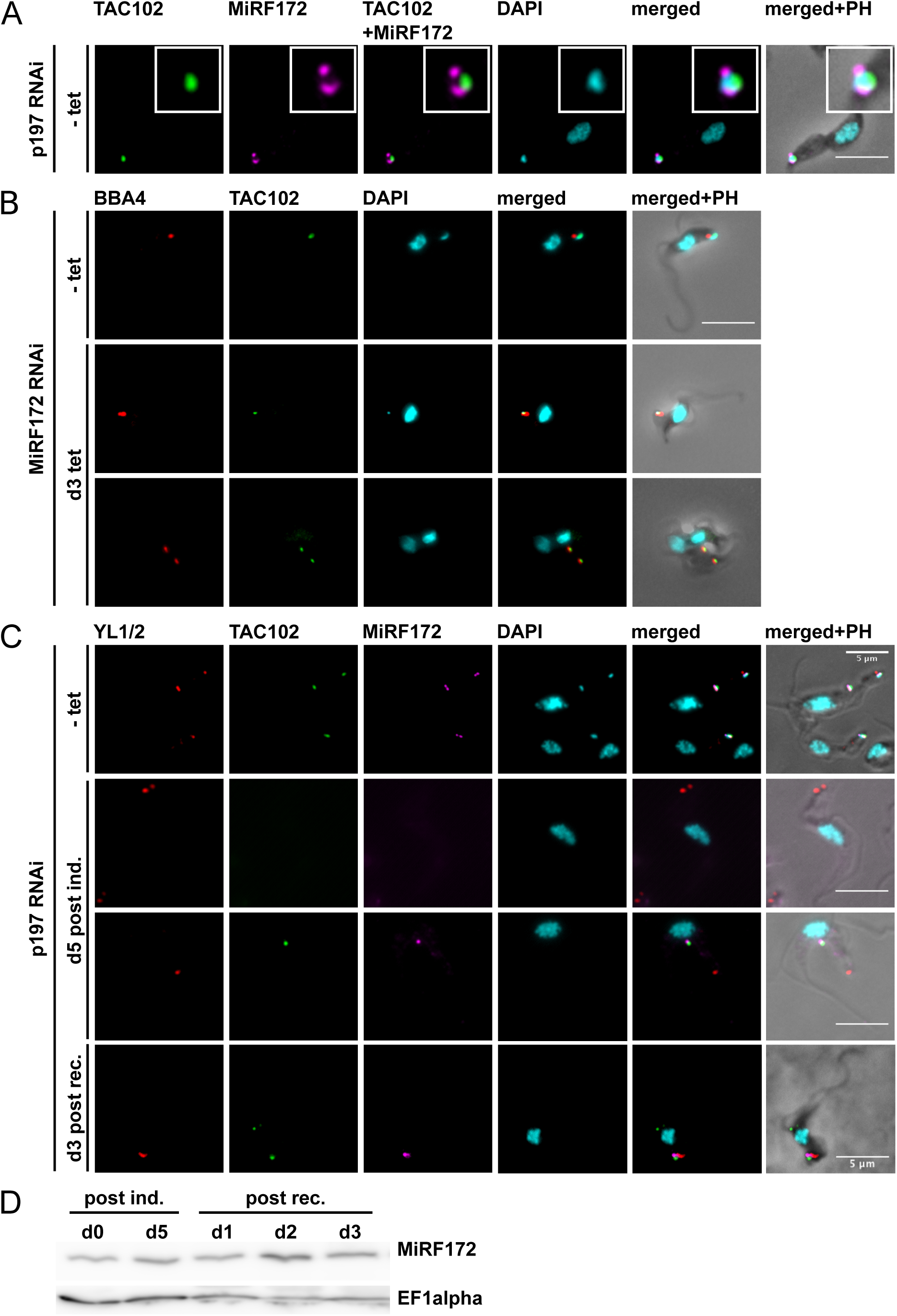
Localization of MiRF172 and TAC102 in cells without kDNA. **A)** Colocalization of MiRF172-PTP with TAC102 in *γ*L262P p197RNAi BSF cells. Localization of MiRF172-PTP (magenta), TAC102 (green) represented by maximum intensity projections from immunofluorescence microscopy image stacks of *γ*L262P p197RNAi BSF *T. brucei* cells. MiRF172-PTP was detected by the α-Protein A antibody. TAC102 was detected with the anti-TAC102 monoclonal mouse antibody (green). The kDNA and the nucleus were stained with DAPI (cyan). **B)** Localization of TAC102 (green) in MiRF172 RNAi BSF cells. The pictures were obtained under the same conditions as described in A. The basal bodies (red) were detected by the monoclonal antibody BBA4. **C)** Colocalization of MiRF172-PTP (magenta) with TAC102 (green) and basal bodies (red) in *γ*L262P p197RNAi BSF cells. The pictures were obtained by same conditions as in A. The basal bodies were detected by the YL1/2 monoclonal antibody. **D)** Western blot analysis of *γ*L262P p197RNAi BSF cells. Total protein isolated from uninduced cells (d0), cells induced with tetracycline for five days (d5) and cells released from p197 RNAi at day 1, 2 and 3 post-recovery were used. C-terminal PTP tagged MiRF172 was detected by a PAP antibody. EF1alpha serves as a loading control. Scale bars = 5 μm

### TAC is required for proper MiRF172 localization

We then wondered if the TAC structure itself has an impact on the localization of MiRF172. For this we created a cell line that allows inducible depletion of p197, a TAC component in the EZF which leads to disruption of the TAC connection between the basal bodies and the kDNA and mislocalization of TAC102 (Gheiratmand et al., 2013; Hoffmann et al., 2017). After five days of p197 mRNA depletion >98% of the cells had lost the kDNA as described previously (Hoffmann et al., 2017) and approximately half of the cells had no signal for MiRF172 as well as TAC102 (Fig. 6C), while the other half of the cells showed a signal for MiRF172 and TAC102 both in close proximity but not at the proper wild type position in the mitochondrion (Fig. 6C). Three days after recovery from p197 RNAi the TAC102 protein was relocalizing properly in vicinity of the basal body as previously described (Hoffmann et al., 2017). For MiRF172 we found a similar behavior. After the recovery from p197 RNAi MiRF172 relocalized in the proximity of TAC102 however it mostly remained as a single spot, rather than two spots as observed in the wild type situation (Fig. 6C).

## Discussion

MiRF172 has no similarities to any other proteins except for the low complexity regions of the C-terminus (Fig. 1). Here, some weak similarities to trfA, a general transcription corepressor and a putative kinase both from Dictyostelium can be found (Aslett et al., 2009). Consistent with its essential function in kDNA replication, MiRF172 is conserved in the currently sequenced Kinetoplastea (Fig. 1A). Of the two recognizable domains (Fig. 1B) the C-terminal alanine/lysine rich region is present in all currently sequenced Kinetoplastea, while the poly-Q stretch is only found in the genus of trypanosomes. We speculate that this variation in the MiRF172 sequence is related to the two kDNA replication models that essentially differ in the reattachment process, which have been proposed for Crithidia and *T. brucei* (Jensen and Englund, 2012). MiRF172 localizes to two regions around the mitochondrial genome that have been described as the antipodal sites in numerous publications (Jensen and Englund, 2012; Povelones, 2014), however the actual composition and dynamics during the kDNA replication cycle as well as the relative position of the individual components in this large structure remain mostly unknown. MiRF172 is present throughout the entire mitochondrial replication and segregation phase in *T. brucei* similar to the primases PRI1 and PRI2, the helicases Pif1 and Pif5, the endonuclease SSE1 and the mitochondrial type II topoisomerase (Engel and Ray, 1999; Hines and Ray, 2010; Hines and Ray, 2011; Li et al., 2007; Liu et al., 2006; Liu et al., 2010). This is different from the polymerase Pol ß, for example that is only at the antipodal sites during replication (Bruhn et al., 2010; Saxowsky et al., 2003). We describe MiRF172 as a novel mitochondrial genome replication factor in *T. brucei*. The current model predicts that replication of the minicircles is initiated after the release into the KFZ by a topoisomerase activity, through binding of the UMSBP and several replication factors, including the two polymerases PolIB and PolIC (Bruhn et al., 2010; Bruhn et al., 2011; Milman et al., 2007). The minicircles are then moved to the antipodal sites by an unknown mechanism. At the antipodal sites primer removal by a single strand endonuclease SSE-1 and the helicase Pif5 are initiated after which gap filling by polymerase Pol ß and finally sealing of most of the gaps through ligase LIG kß occur (Downey et al., 2005; Engel and Ray, 1999; Klingbeil et al., 2002; Saxowsky et al., 2003). Afterwards the minicircles are reattached to the growing network, likely by topoisomerase activity (Wang and Englund, 2001). In the kDNA disc the last minicircle gaps are repaired through a combination of Polß-Pak and the DNA ligase LIG kα and likely other proteins (Downey et al., 2005; Klingbeil and Englund, 2004; Klingbeil et al., 2002). Based on the current model, an accumulation of gapped free minicircles as detected in the MiRF172 RNAi analysis, points towards a function of MiRF172 in the reattachment process. The only other currently known protein to be involved in reattachment is the mitochondrial topoisomerase TOPOII, which is also localized at the antipodal sites. Thus, we predict MiRF172 and TOPOII to interact with each other in the process of minicircle reattachment in *T. brucei*. One could imagine that MiRF172 might aid the topoisomerase in the discrimination between replicated and non-replicated minicircles. In the future, biochemical co-immunoprecipitation studies should allow us to test this model.

The proximity of the kDNA replication and segregation machinery in *T. brucei* could suggest a physical interaction between the two processes in the KFZ. This is supported by the biochemical fractionations indicating that MiRF172 remains associated with the isolated flagellum and is present in the pellet fraction of the digitonin extraction even after DNAseI treatment (Fig. S2).

Interestingly MiRF172 has no impact on proper TAC biogenesis, however TAC biogenesis is required for proper MiRF172 localization since loss of the TAC structure also leads to loss of proper MiRF172 localization. Even in the absence of kDNA in the “petite” trypanosomes, MiRF172 localizes close to TAC102 although in a different conformation. This suggests that the TAC does provide important localization information for replication proteins such as MiRF172. It will be interesting to test if this is also true for the topoisomerase and other antipodal site proteins.

## Material and Methods

### *T. brucei* cell culture conditions

Monomorphic *T. brucei* BSF NYsm (Wirtz et al., 1999) and NYsm-derived γL262P (Dean et al., 2013) cells were cultured in Hirumi-modified Iscove’s medium 9 (HMI-9) supplemented with 10% fetal calf serum (FCS) and incubated at 37°C and 5% CO_2_. Procyclic form (PCF) 427 *T. brucei* cells were cultured in semi-defined medium-79 (SDM-79) supplemented with 10% FCS at 27°C. Depending on the cell line 5 μg/ml blasticidin, 2.5 μg/ml geneticin, 2.5 μg/ml hygromycin, 2.5 μg/ml phleomycin or 0.5 μg/ml puromycin were added to the medium. Expression of the RNAi construct was induced through the addition of 1 μg/ml tetracycline. NYsm BSF, 427 PCF trypanosomes were obtained from the established collection of the Institute of Cell Biology, University of Bern, Bern, Switzerland. The γL262P strain of BSF cells is a kind gift of A. Schnaufer.

### Transfections of *T. brucei* cells

For transfections, 10 μg of linearized plasmid or PCR product were dissolved in 100 μl BSF transfection buffer (90 mM Na-phosphate pH 7.3, 5 mM KCl, 0.15 mM CaCl_2_, 50 mM HEPES pH 7.3) (Burkard et al., 2007). 4×10^7^ mid-log phase BSF cells were pelletized and resuspended in 100 μl BSF transfection buffer containing the DNA. The cells were transferred into Amaxa Nucleofector cuvettes and transfections were conducted in the Amaxa Nucleofector II using program Z-001 (panel V 1.2 kV, panel T 2.5 kV, panel R 186 Ohm, panel C 25 μF). For transfections of PCF cells, 10 μg of PCR product were dissolved in 400 μl Zimmerman postfusion media (ZPFM, 132 mM NaCl, 8 mM KCl, 8mM Na_2_HPO_4_, 1.5 mM KH_2_PO_4_, 0.5 mM MgAc_2_, 0.09 mM CaAc_2_). 10^8^ mid-log phase PCF cells were pelletized and resuspended in the ZPFM containing the DNA and transferred into Amaxa Nucleofector cuvettes. Transfections were conducted with 1500V, 180 ohms, 25μF (BTX). The transfected cells were left to recover for 20 h. Cells were then selected with appropriate antibiotics for correct integration of the construct.

### DNA constructs

The MiRF172 RNAi constructs were targeted against the 4409 to 4719 bp and of the ORF and 1 to 10 bp of the 3’ UTR of the gene Tb927.3.2050. Briefly, a PCR fragment with adaptor sequences was amplified from genomic DNA of NYsm BSF cells, and cloned in two steps into the pTrypRNAiGate vector by Gateway cloning (Kalidas et al., 2011). The final plasmids linearized with NotI HF (NEB) were used for transfection as described above. Expression was induced by addition of 1 μg/ml tetracycline. For C-terminal PTP-tagging of MiRF172, the ORF positions 4404 to 4895 were amplified from genomic DNA of NYsm BSF cells and cloned between ApaI / NotI sites of the pLEW100 based PTP tagging vector (Schimanski et al., 2005). The resulting plasmid was linearized with BsmI prior to transfection. For the C-terminal triple HA-tagging, a PCR with primers containing overhangs complementary to the ORF from 4617 to 4716 and the 3’ UTR from 1 to 99 was performed. The pMOTagging plasmid served as a template (Oberholzer et al., 2006). Both tagging constructs were recombined into the endogenous locus to substitute for one of the Tb927.3.2050 alleles and thus was constantly expressed.

### Immunofluorescence analysis

Approximately 10^6^ cells were spread onto a slide and fixed for 4 min with 4% PFA in PBS (137 mM NaCl, 2.7 mM KCl, 10 mM Na_2_HPO_4_, 2 mM KH_2_PO_4_, pH 7.4). After fixation and washing with PBS cells were permeabilized with 0.2% Triton-X 100 in PBS for 5 min. Then cells were blocked with 4% BSA in PBS for 30 min. After blocking, slides were incubated for 45 or 60 min with the primary antibody followed by washing with PBS + 0.1% Tween-20 (PBST) and incubation with the secondary antibody for 45 or 60 min followed by washing with PBST and PBS. All incubations were performed at room temperature. Primary and secondary antibodies were diluted in PBS + 4% BSA as follows: rat-α-HA (Sigma) 1:1000, polyclonal rabbit-α-Protein-A (Sigma) detecting the PTP epitope 1:2000, mouse BBA4 antibody (Woods et al., 1989) 1:100, rat YL1/2 antibody detecting tyrosinated tubulin as present in the basal body (Kilmartin et al., 1982) 1:100000, Alexa Fluor® 594 Goat-α-Rabbit IgG (H+L) (Life technologies), Alexa Fluor® 488 Goat-α-Rabbit IgG (H+L) (Invitrogen), Alexa Fluor® 488 Goat-α-Rat IgG (H+L) (Life technologies), Alexa Fluor® 488 Goat-α-Mouse IgG (H+L) (Invitrogen), Alexa Fluor® 594 Goat-α-Mouse IgG (H+L) (Molecular probes), Alexa Fluor® 647 Goat-α-Rat IgG (H+L) (Life technologies) all 1:1000. Cells were mounted with ProLong® Gold Antifade Mounting Medium with DAPI (Molecular Probes) and cover slips were added. Images were acquired with the Leica DM5500 B microscope (Leica Microsystems) with a 100× oil immersion phase contrast objective. Images were analyzed using LAS X software (Leica Microsystems) and ImageJ. Significance of the quantification of relative occurrence of kDNA and nucleus in different cell cycle stages was calculated using the two-tailed unpaired t-test.

### Super resolution 3D STED (Stimulated Emission Depletion) microscopy

MiRF172-PTP BSF cells were spread and fixation, permeabilization and blocking were performed as described above. Polyclonal rabbit-α-Protein-A antibody (Sigma) and the Alexa Fluor® 594 goat-α-Rabbit IgG (H+L) antibody were used as described above. Cover glasses (Nr. 1.5) suitable for 3D STED microscopy (Marienfeld) were used. Cells were mounted in ProLong with DAPI as described above. Images were acquired using the SP8 STED microscope (Leica, with a 100× oil immersion objective and the LAS X Leica software) as z-stacks with a z-step size of 120 nm. For the MiRF172-PTP signal the 594 nm excitation laser, the 770 nm depletion laser were used. The DAPI signal was acquired with confocal settings. Images were deconvoluted with the Huygens professional software.

### Transmission electron microscopy (TEM)

Embedding of the cells and thin sectioning for TEM was performed as described previously (Trikin et al., 2016). Images of the thin sections were obtained by the FEI Morgani electron microscope (Tungsten cathode). The microscope was equipped with a digital camera (Morada, 12 megapixel, Soft Imaging System) and the AnalySIS iTEM image analysis software. The kDNA structure of uninduced cells (-tet) and induced cells at day three upon induction (+tet d3) was measured using ImageJ. All images are taken at a magnification of 28000×. Significance of the results was calculated using the two-tailed unpaired t-test.

### SDS-PAGE and western blotting

Whole cell lysates were used for western blot analysis. Cells were washed in PBS and resuspended in 1× Laemmli buffer (12 mM Tris-Cl pH 6.8, 0.4% SDS, 2% glycerol, 1% β-mercaptoethanol, 0.002% bromophenol blue) in PBS and heated for 5 min at 95 °C. Approximately 5×10^6^ cells were loaded onto a 4% or 6% gel, resolved and then blotted (BioRAD blotting system) onto PVDF Immobilon®-FL transfer membranes (0.45 μm, MILLIPORE) for 1 h at 100 V. Membranes were blocked in PBST + 5% skim milk powder. The rabbit peroxidase anti-peroxidase soluble complex (PAP) was diluted 1:2000 in PBST + 5% skim milk and incubated for 30 min at RT. The mouse-anti-EF1alpha (Santa Cruz), rat-anti-HA and the rabbit-anti-HA antibodies (Sigma) were used 1:1000 in PBST + 5% skim milk. Secondary antibodies were: swine anti-rabbit HRP-conjugate (1:10000, Dako) and rabbit anti-rat HRP-conjugate (1:10000, Dako), all in PBST + 5% skim milk. After each incubation with the antibodies, the membranes were washed 3× for 5 min in PBST and 1× for 5 min in PBS. SuperSignal West Femto Maximum Sensitivity Substrate (Thermo Scientific) and Amersham^TM^ Imager 600 (GE Healthcare Life Sciences) were used to visualize the protein bands on the blots.

### Northern blotting

Total RNA was extracted from mid-log phase MiRF172 RNAi BSF cells with 1 ml of RiboZol ™ (Amresco) per 5×10^7^ cells. For northern blot analysis, 10 μg of total RNA was separated for two hours at 100 V in a 1% agarose gel containing 6% formaldehyde. RNA was blotted onto Hybond nylon membranes with 20× SSC (3 M NaCl, 0.3 M Na-citrate pH 7) by capillary transfer. The RNA on the nylon membrane was cross-linked with Stratagene UV-Stratalinker. Membranes were pre-hybridized at 65°C for one hour in hybridization solution (5× SSC, 1:12.5 100× Denhardt’s (2% BSA, 2% polyvinylpyrrolidone, 2% Ficoll), 50 mM NaHPO_4_ pH 6.8, 1% SDS, 100 μg/ml salmon sperm DNA). The sequence specific probe for MiRF172 mRNA was generated by PCR (Primers used: 5’-ggggacaagtttgtacaaaaaagcaggctCCCTGAGAAGGAACTTGAGC-3’ and 5’-ggggaccactttgtacaagaaagctgggtGGCTGCTCATCTACCGCTT-3’). The probes were denatured in ddH_2_O for 5 min at 95°C. Random primed DNA labeling kit (Roche) was used according to the manufacturer’s manual to label the probes. For normalization, 1.8 μl 18S rRNA probes (10 μM, (Trikin et al., 2016)) was mixed with 12.5 μl H_2_O, 2.7 μl gamma-^32^P-ATP (1 MBq), 2 μl PNK buffer (10×) and 1 μl T4 PNK and incubated for 30 min at 37°C. Reactions were stopped with 5 μl EDTA (0.2 M) and 75 μl TE buffer (1×) and incubated 5 min at 95° C. Probes were quenched for 2 min on ice and 50 μl of probe mixture was added to the membrane. Probes and pre-hybridized membranes were incubated over night at 65°C and washed in 2× SSC, 0.1% SDS and/or 0.2× SSC, 0.1% SDS at 60°C. Blots probed for MiRF172 RNA were exposed for 24 hours, when probed for 18S rRNA - for approximately 15 min, to storage phosphor screens in metal cassettes (Amersham Bioscience) and scanned by a Storm PhosphoImager (Amersham Bioscience). ImageJ was used for image analysis and quantification.

### Southern blotting

Total DNA was isolated from mid-log phase MiRF172 RNAi BSF cells. For this, cells were washed in PBS and resuspended in 1ml of phenol per 5×10^7^ cells. Experimental procedure and analysis was performed as described previously (Trikin et al., 2016). 5 μg of total DNA either undigested (for detection of free minicircles) or digested with HindIII and XbaI (for detection of total mini- and maxicircles) was resolved in 1% agarose gel within 35 min for total minicircles or 2 h for free minicircles at 135 V in 0.5× TAE buffer. Sequence specific probes for minicircles were generated from a PCR fragment (approx. 100 bp of the conserved minicircle sequence (Trikin et al., 2016)) amplified from total DNA of NYsm BSF *T. brucei*. The maxicircle probe (Trikin et al., 2016) was amplified from total DNA of NYsm BSF *T. brucei* too using the following primers: 5’ - CTAACATACCCACATAAGACAG-3’ and 5’ -ACACGACTCAATCAAAGCC-3’ (Liu et al., 2006). For the normalization, a tubulin probe (binding to the intergenic region between α- and β-tubulin, fragment size 3.6 kb, (Trikin et al., 2016)) was used and it was generated and labelled in the same way as the minicircle and maxicircle probes. Blots were exposed for 24, 48 or 72 hours to storage phosphor screens in metal cassettes (Amersham Bioscience) and scanned by Storm PhosphoImager (Amersham Bioscience). ImageJ was used for image analysis and quantification. Significance of the results was calculated using the two-tailed unpaired t-test.

### Flagellar extraction

For flagellar extraction, EDTA was added to BSF cells in medium with an end concentration of 5 mM. Cells were washed with PBS and then resuspended in extraction buffer (10 mM NaH_2_PO_4_, 150 mM NaCl, 1 mM MgCl_2_) containing 0.5% TritonX-100, on ice. After one washing step with extraction buffer, cells were incubated on ice for 45 min in extraction buffer containing 1 mM CaCl_2_ and then subjected to immunofluorescence analysis as described above.

### Digitonin fractionations

For digitonin fractionation 10^7^ cells were collected and washed with PBS. Then they were resuspended in SoTE buffer (0.6 M sorbitol, 2 mM EDTA, 20 mM Tris-HCl, pH 7.5). Digitonin was added to a final concentration of 0.025% or 1% and the mixture was incubated on ice for 5min. To separate the fractions, cells were centrifuged at 8000 rcf for 5 min at 4°C. Both fractions (supernatant and pellet) were mixed with Laemmli buffer for western blot analysis.

## Acknowledgement

For the BBA4 and YL1/2 antibodies we would like to thank Keith Gull. We acknowledge Bernd Schimanski and Beat Haenni for technical assistance, Anneliese Hoffmann for critical reading of the manuscript, Borka Jojic and Roman Trikin for critical discussion. Electron microscopy sample preparation and imaging was performed with devices supported by the Microscopy Imaging Center (MIC) of the University of Bern, Switzerland. For financial support, we thank the Berne University Research Foundation and the Novartis Foundation.

## Competing Interest

The authors declare no competing financial, personal or professional competing interests.

## Author Contribution

Torsten Ochsenreiter (TO) and Simona Amodeo (SA) designed the experiments. SA performed the experiments and analyses. Martin Jakob (MJ) and SA designed the model of MiRF172 localization. TO and SA wrote the manuscript. MJ read and participated in the corrections of the initial draft.

